# C-terminal disulfide bond in SUN2 regulate dynamic remodeling of LINC complexes at the nuclear envelope

**DOI:** 10.1101/2022.06.08.495410

**Authors:** Rahul Sharma, Martin W. Hetzer

## Abstract

The LINC complex tethers the cell nucleus to the cytoskeleton to regulate mechanical forces during cell migration, differentiation, and diseases. The function of LINC complexes relies on the interaction between highly conserved SUN and KASH proteins that form higher-order assemblies capable of load bearing. These structural details have emerged from *in vitro* assembled LINC complexes however, the principles of *in vivo* assembly remain obscure. Here, we report a conformation-specific SUN2 antibody as a tool to visualize LINC complex dynamics *in situ*. Using imaging, biochemical and cellular methods, we find that conserved cysteines in SUN2 undergo KASH dependent inter- and intra- molecular disulfide bond rearrangements. Disruption of the SUN2 terminal disulfide bond compromises SUN2 localization, turnover, LINC complex assembly as well as cytoskeletal organization and cell migration. Moreover, using pharmacological and genetic interventions, we identify components of the ER lumen as SUN2 cysteines redox state regulators. Overall, we provide evidence for SUN2 disulfide bond rearrangement as a physiologically relevant structural modification that regulates LINC complex functions.

## Introduction

Apart from harboring and organizing the genome, the cell nucleus plays a crucial role in integrating intra- and extra-cellular mechanical cues to generate an appropriate cellular response. The Linker of Cytoskeleton and Nucleoskeleton (LINC) is a specialized protein complex that forms the molecular basis of this mechanical signaling by physically connecting the cytoskeleton to the nuclear lamina (NL). The LINC complex is comprised of a highly conserved Sad1/UNC-84 (SUN) domain containing protein and Klarsicht/ANC-1/Syne-1 homology (KASH) domain containing proteins that traverse the inner and outer nuclear membranes (INM/ONM), respectively. SUN and KASH proteins directly interact in the nuclear envelope (NE) lumen (i.e. perinuclear space), which is continuous with the endoplasmic reticulum (ER) lumen. Additionally, SUN and KASH proteins interact with the NL and Actin/Microtubule network to physically tether the nucleus to the rest of the cell(Crisp et al., 2006) (Fig 1A). The LINC complex is important for key cellular processes like cell migration, nuclear positioning and anchorage, cell differentiation, and mechanotransduction(Zhang et al., 2009, 2007; Gant et al., 2010; Lombardi et al., 2011; Carley et al., 2021; Cain et al., 2018; Déjardin et al., 2020). Moreover, since SUN and KASH proteins are associated with muscular dystrophy, cardiomyopathy and cancer(Chen et al., 2012; Meinke et al., 2014; Stewart et al., 2019; Matsumoto et al., 2015; Maurer and Lammerding, 2019; Battey et al., 2020), the understanding of the molecular basis of LINC complex assembly and function could lead to future therapeutics.

**Figure 1:**
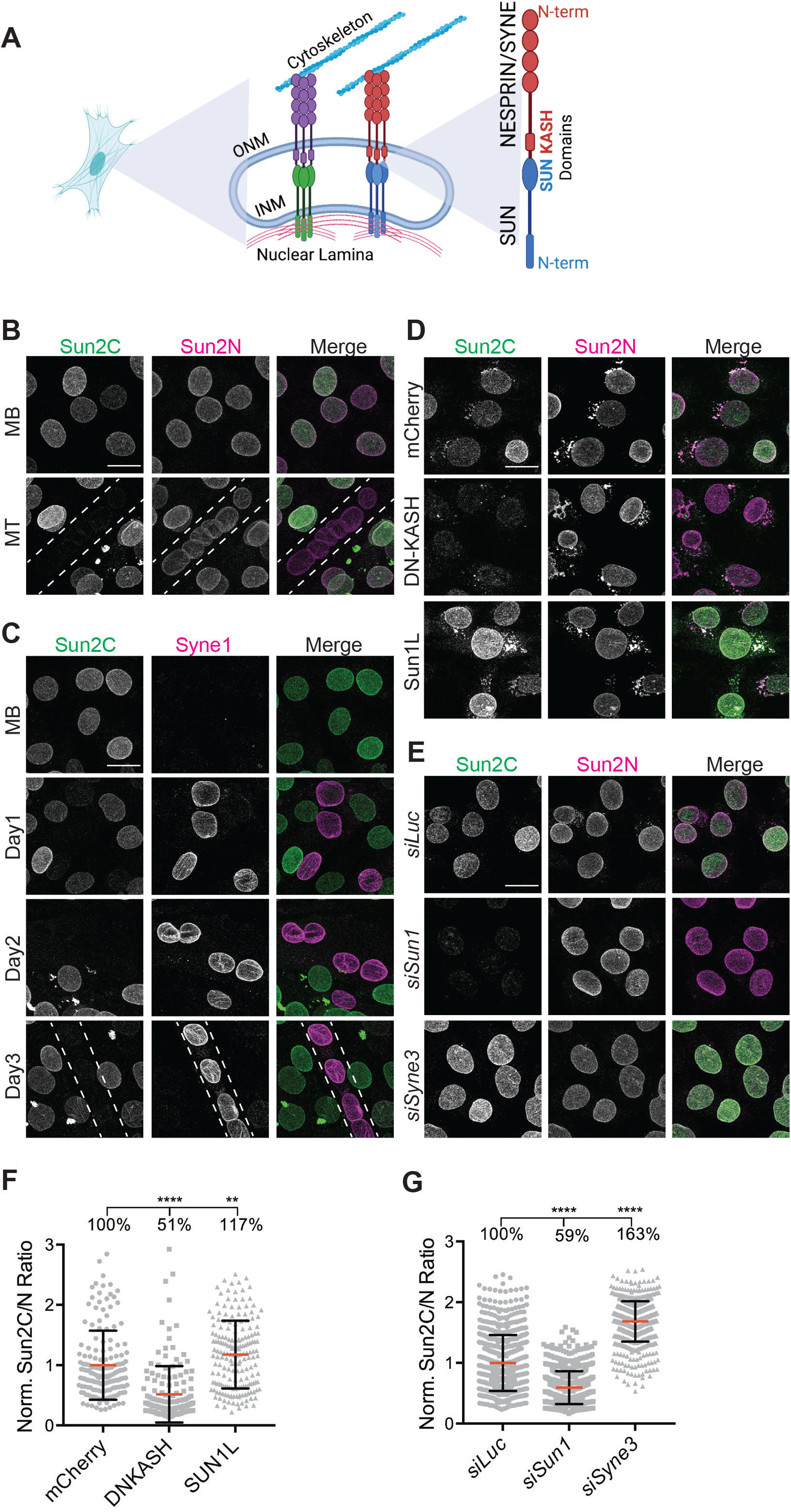
Nesprin interaction masks the epitope of a monoclonal SUN2 antibody. (A) Schematic representation of the LINC complex at the nuclear envelope. (B) Representative images show Sun2C and Sun2N antibody staining in C2C12 Myoblast cells (MB) or Myotubes (MT). Dotted line represents MT outline. (C) Representative images of C2C12 MB at different stages of differentiation (Day1-3) into MTs costained for SUN2 and NESPRIN1/SYNE1. (D). Representative images of C2C12 MB cells stained with Sun2C and Sun2N Ab, 24hr post-transfection with mCherry, DN-KASH and SUN1L constructs. (E) Representative images of C2C12 MB cells 72hr post-transfection with siRNAs against *Luciferase, Sun1 or Nesprin3/Syne3*, co-stained with Sun2C and Sun2N antibody. (F and G) Quantitation for images in (D) and (E) respectively. Scatter plot shows the ratio of total nuclear fluorescence intensity of Sun2C over Sun2N for individual cells in different conditions and normalized to average ratio of mCherry (for F) or *siLuc* (for G). Mean (Red bar) and SD (black bar) are represented. Data pooled from two independent experiments. n>100 (for E) and n>600 (for F) cells per condition. *t-test* was applied to calculate statistical significance against mCherry (for F) and *siLuc* (for G). * p<0.05, ** p<0.01, **** p<0.0001. Values show mean percentage of that group. All images are max intensity projections of confocal z-stacks. All scalebars are 20um.

Mammals express five different SUN-containing proteins of which, SUN1 and SUN2 are ubiquitous and partially redundant(Kai et al., 2009). Apart from the conserved SUN domain, SUN proteins harbor additional functional domains: (i) a lamin binding domain that interacts with A/B-type lamins and helps to anchor the protein in the INM(Crisp et al., 2006), (ii) two coiled coil domains (CC1, CC2) that reside in the NE lumen and seem to regulate protein conformation(Nie et al., 2016). In addition, five different KASH-containing proteins (NESPRIN/SYNE) and multiple isoforms have been reported in mammals generating a remarkable diversity among the KASH family(Zhang et al., 2005). NESPRINS are megadalton proteins comprising of cytoskeleton-interacting domain, spectrin repeat region and a small 20-30 amino acid KASH domain (Meinke and Schirmer, 2015)(Fig 1A). Although *in vitro* data suggests promiscuous interaction among different SUN and NESPRIN proteins(Kim et al., 2015), there is evidence for specialized SUN:NESPRIN pairs. For example, SUN1:NESPRIN1 and SUN2:NESPRIN2 pairs are required for forward and rearward nuclear movement in migrating cells, respectively(Zhu et al., 2017). How endogenous SUN proteins choose NESPRIN partners remains unclear. Moreover, acquisition of new forms of SUN and NESPRIN proteins in vertebrates, points towards an underexplored functional diversity and emergence of LINC independent functions over the course of evolution. Currently there are no tools or techniques available to distinguish between bound and unbound SUN/KASH proteins *in vivo*.

X-Ray crystallography of *in vitro* assembled truncated fragments of SUN and KASH proteins have been invaluable in providing insights into SUN-KASH interaction and LINC complex assembly. The emerging model describes a heterohexameric complex wherein SUN homotrimer interacts with three KASH peptides(Sosa et al., 2012; Wang et al., 2012). SUN oligomerization has been shown as a prerequisite for KASH interaction but the principles behind SUN oligomerization are unclear with conflicting models(Jahed et al., 2021). One study suggests that an interplay between an activator CC1 and inhibitor CC2 domain regulates SUN2 monomer to trimer ratios and thereby, SUN-KASH interactions(Nie et al., 2016). Another recent study proposes a self-locking state wherein two SUN trimers are locked head-to-head and unlock at the INM for KASH interaction(Cruz et al., 2020). Conversely, an experimentally validated intermolecular disulfide bridge between cysteine residues in SUN and KASH domains has been described that is dispensable for KASH binding but required for maximal force transmission(Cain et al., 2018; Jahed et al., 2015). Lastly, two conserved C-terminal cysteines in SUN2 have been predicted to form an intramolecular disulfide bond(Sosa et al., 2012). However, the importance of these cysteines remains to be evaluated. An obvious limitation of these structural studies is the use of truncated fragments of proteins in non-physiological conditions. Thus, several key questions remain unanswered: Do these autoinhibitory conformations exist *in vivo?* How do SUN proteins remain inactive and KASH unbound state in ER but activate after reaching INM? What controls SUN activation step? How do LINC complexes respond to dynamic changes in mechanical forces? Lack of appropriate tools remains a major limitation to investigate these questions.

Here, we have identified a key structural change in SUN2 associated with assembly and disassembly of LINC complex *in situ*. We report the discovery of a conformation-specific SUN2 antibody that specifically recognizes a SUN2 fraction that is not bound to KASH. Using a semiquantitative imaging assay, we show that KASH bound and unbound SUN2 fraction changes dramatically during cell proliferation, differentiation and migration. Molecular dissection of the antibody epitope, revealed a key structural feature of SUN-KASH interaction, namely that KASH interaction oxidizes the highly conserved SUN2 cysteine residues thereby, masking the epitope. Moreover, genetic and pharmacological perturbation experiments, provide evidence for the functional conservation of SUN2 cysteine residues and identify potential redox regulators present in the ER. Overall, we provide a structure-function analysis for SUN2 proteins in physiological context.

## Results

### Discovery of a conformation-specific SUN2 antibody

Muscle differentiation changes the overall NE proteome and additionally alters the stoichiometry of LINC complex proteins(Chen et al., 2006; Randles et al., 2010; Loo et al., 2019; Wilkie et al., 2011). To understand changes in these proteins during myogenesis, we performed immunofluorescence (IF) staining in C2C12 myoblasts (MB) and terminally differentiated myotubes (MT) using two highly specific monoclonal antibodies against the C-terminus (Sun2C) and N-terminus (Sun2N) of SUN2 which we validated by siRNA knockdowns (Supp Fig 1A-C). Interestingly, the two antibodies behaved differently. The antibody (Sun2N) targeting the N-terminus epitope of SUN2, resulted in homogenous nuclear rim staining confirming the presence of SUN2 in both MBs and MTs (Fig 1B, Sun2N). In contrast, we observed a striking heterogeneity of nuclear staining in MBs and failed to obtain a Sun2C signal in MTs altogether (Fig 1B, Sun2C). These results suggest that the epitope of Sun2C is masked in MT.

Onset of myogenesis is marked by an upregulation of a MT specific NESPRIN1/SYNE1 that is required for muscle differentiation and fundamentally alters LINC complex composition in MT(Stroud et al., 2017; Espigat-Georger et al., 2016). Therefore, we tested whether gain of SYNE1 is associated with the loss of Sun2C epitope in MTs. Upon co-stained MBs at different stages of differentiation, we found that the loss of Sun2C epitope coincided with SYNE1 upregulation as early as Day1 and remained inaccessible thereafter (Fig 1C). Since SYNE1 is a binding partner of SUN proteins, this suggested a link between NESPRIN upregulation and epitope masking of Sun2C antibody in MTs.

Differentiation changes LINC complex composition, but whether the masking of Sun2C epitope in MTs is dependent on differentiation or represents an independent change in LINC complex, remained unclear. To address this question, we mimicked NESPRIN upregulation in undifferentiated MBs by transiently expressing a dominant negative form of NESPRIN (DN-KASH) that lacks the actin binding domain but retains the KASH domain(Stewart-Hutchinson et al., 2008; Lombardi et al., 2011). Upon co-staining with both SUN2 antibodies, we found that Sun2C failed to recognize SUN2 in the presence of DN-KASH compared to mCherry alone expressing cells (Fig 1D) with significantly reduced Sun2C intensity (Fig 1F, see Methods for quantitation). This shows that Sun2C epitope masking is independent of differentiation but dependent on SUN2:DN-KASH interaction. We hypothesized that, if Sun2C epitope is masked in KASH bound SUN2, then conversely, Sun2C epitope should be unmasked in KASH unbound SUN2. To test this, we transiently expressed SUN1L, a previously reported luminal version of SUN1 that outcompetes endogenous SUN proteins for KASH binding(Stewart-Hutchinson et al., 2008). As expected, cells expressing SUN1L showed enhanced binding of Sun2C and significantly increased intensities (Fig 1D, 1F). Overall, this experiment shows that Sun2C epitope is masked in a SUN2:KASH complex but remains accessible for unbound SUN2.

SUN1 and SUN2 both compete for limited KASH peptides at the NE. We hypothesized that in the absence of SUN1, a majority of SUN2 would bind KASH and similarly, loss of NESPRIN would shift the balance to more unbound SUN proteins. To confirm that Sun2C Ab responds to changes in SUN2:KASH interactions, we transiently depleted either SUN1 or NESPRIN3/SYNE3 in MBs (Supp Fig 1D-F) and co-stained with SUN2 antibodies. As expected, depleting SUN1, lead to a dramatic loss and depletion of NESPRIN3 increased Sun2C intensity (Fig 1E, 1G). Taken together, these data demonstrate that Sun2C is a conformation-specific SUN2 antibody that only recognizes unbound SUN2 molecules and by using this antibody we show that SUN2 exists predominantly in a KASH bound state in MTs.

### KASH interaction alters intermolecular disulfide bonds in SUN2

KASH dependent Sun2C epitope masking presents a rare and interesting opportunity to investigate LINC complex assembly and dynamics in a physiological context. *In vitro* structure studies have revealed SUN2 oligomerization as a prerequisite for KASH engagement(Sosa et al., 2012). Therefore, we wondered whether the progressive loss of Sun2C signal observed during myogenesis correlates with changes in SUN2 oligomerization. To test this, we biochemically resolved SUN2 oligomers from cells undergoing differentiation, using a previously published non-reducing PAGE method (Lu et al., 2008). We found that undifferentiated MBs contained SUN2 monomers (Sun2(M)) and higher molecular weight oligomers (Sun2(O)). In contrast, fully differentiated Day6 MTs lost the majority of SUN2 oligomers (Fig 2A -DTT, 2D). Additionally, all oligomers were lost upon treatment with a reducing agent, Dithiothreitol (DTT) (Fig 2A +DTT). Since non-reducing gels can only resolve disulfide linked species, we conclude that SUN2 undergoes an intermolecular disulfide bond rearrangement during C2C12 differentiation.

**Figure 2:**
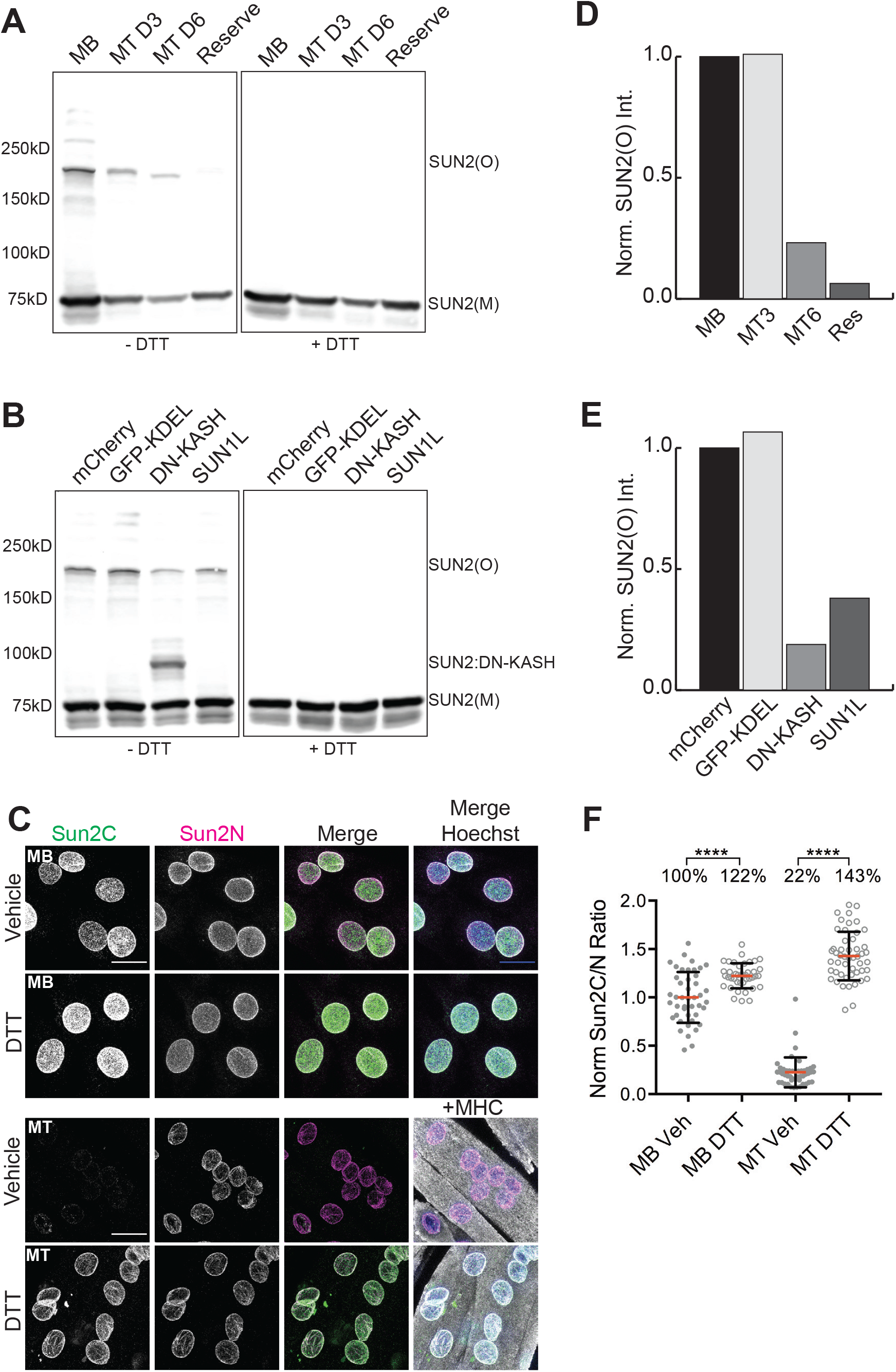
Nesprin interaction changes SUN2 disulfide linkage. (A) Western blot shows differential migration of SUN2 monomer (SUN2(M)) and oligomer (SUN2(O)) bands in C2C12 Myoblast (MB), Day3 and Day6 Myotubes (MT D3, D6) and Reserve cells in non-reduced (left panel, -DTT) or reduced (right panel, +DTT) conditions. (B) Similar to (A) Lysates from MBs, 24hr post-transfection of mCherry, GFP-KDEL, DN-KASH or SUN1L constructs. A separate band for SUN2:DN-KASH complex is observed at ~100kD. (C) Representative images of MB or MTs treated with Vector (Water) or 5uM DTT and co-stained with Sun2C and Sun2N antibodies. Myosin heavy chain (MHC) staining was used as a MT marker. (D) and (E) show densitometry analysis of (A) and (B) respectively. Bar plot represents ratio of SUN2(O) to SUN2(M) band intensity per condition; normalized to MB (for D) and mCherry (for E). (F) Quantification for (C) Scatter plot shows the ratio of total nuclear fluorescence intensity of Sun2C over Sun2N signal for individual cells in different conditions and normalized to average ratio of Vector. n>40 cells per condition. Mean (Red bar) and SD (black bar) are represented. *t-test* was applied to calculate statistical significance against corresponding Vehicle control. * p<0.05, ** p<0.01, **** p<0.0001. Values show mean percentage of that group. All images are max intensity projections of confocal z-stacks. Scalebar is 20um.

Previously, it has been shown that SUN2 forms a disulfide bond with NESPRINs to increase the structural integrity of LINC complexes(Sosa et al., 2012; Cain et al., 2018). To test whether KASH interaction leads to loss of SUN2 oligomers, we transfected MBs with previously described DN-KASH, SUN1L, mCherry control constructs and a GFP-KDEL control that expresses GFP in ER lumen similar to SUN1L(Lombardi et al., 2011). Interestingly, in our biochemical analysis, we found that SUN2 oligomers (Sun2(O)) decreased dramatically in the presence of DN-KASH and an additional band representing disulfide linked SUN2:DN-KASH was detected (Fig 2B -DTT, 2E). The SUN2 oligomer band also decreased upon SUN1L expression but remain unaffected in mCherry and KDEL controls (Fig 2B -DTT, 2E). Again, all oligomers were lost under reducing conditions (Fig 2B +DTT). This data suggests that assembly and disassembly of SUN2 LINC complex is associated with changes in SUN2 disulfide linkages. Lastly, to demonstrate that these disulfide rearrangements observed on the gel are physiologically relevant, we treated MBs and MTs with DTT prior to fixation and performed IF with Sun2C and Sun2N Abs, which was sufficient to retrieve the Sun2C epitope in MTs and significantly increased Sun2C intensity compared to control (Fig 2C, 2F). This further confirms that SUN2:KASH interaction leads to formation of disulfide bridges that mask the epitope for Sun2C *in situ*.

### Conserved C-terminal cysteines determine SUN2 NE targeting, protein turnover and Sun2C epitope masking

Since disulfide bonds are formed between cysteine residues, we focused on the amino acid (AA) composition of SUN2 to unravel the principles underlying SUN2:KASH mediated disulfide linkages. Mouse SUN2, which is a 731AA protein, contains only three highly conserved cysteine residues that reside in the SUN domain (Supp Fig 2). Structural data shows that SUN2 Cys577 interacts with Cys-23 in the KASH domain of NESPRINs whereas, Cys615 and Cys719 form an intramolecular disulfide bridge (Fig 3A)(Sosa et al., 2012)(Wang et al., 2012)(Cruz et al., 2020). To understand which cysteines are involved in Sun2C masking, we generated a N-term GFP-tagged SUN2 (WT) and alanine substituted cysteine mutant (C577A and C719A) constructs. Upon stable expression in MBs, we observed that WT and C577A expressing lines showed high GFP expression (Fig 3B GFP panel, 3C). The ectopically expressed protein localized properly to the NE similar to endogenous SUN2 (Fig 3B, 3D). On the contrary, C719A mutant showed comparatively lower expression and accumulated predominantly in the ER (Fig 3B-D).

**Figure 3:**
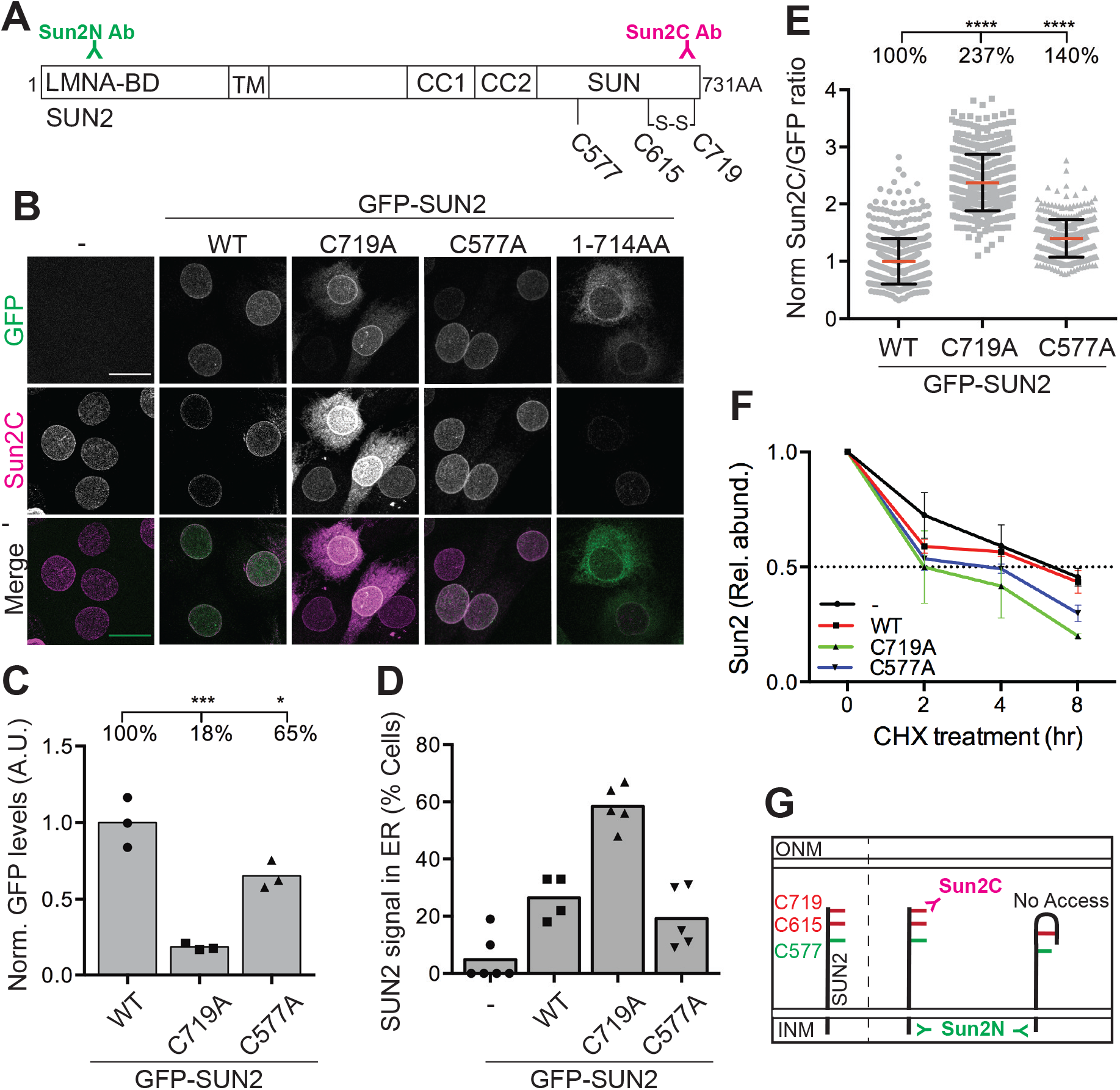
SUN2 terminal disulfide bond is required for proper NE localization and turnover of SUN2. (A) Schematic diagram depicting different domains within mouse SUN2 full length protein (1-731AA): Lamin A Binding region (LMNA-BD), Transmembrane domain (TM), Coiled Coil1 and 2 (CC1 and CC2), SUN domain (SUN). Three Cysteines present in SUN2 are labelled according to their amino acid position in the peptide (C577, C615, C719). S-S represents a disulfide bond. Sun2N and Sun2C represents the corresponding epitopes in SUN2 peptide for their respective antibodies. (B) Representative images of C2C12 cells stably expressing N-terminal GFP-tagged SUN2 (WT) or Alanine substituted Cysteine mutants (C719A and C577A) stained with Sun2C antibody. Additionally, a mutant with 17AA deletion at SUN2 C-terminus (1-714AA) is included that does not recognize Sun2C Ab. (C) Bar graph shows the total level of GFP (normalized to aTubulin) in WT and mutant lines based on densitometry analysis of western blots. Each symbol represents an individual replicate of three independent experiments. *t-test* was performed to measure significance against WT. (D) Bar graph shows percentage of cells with SUN2 signal in ER for different cell lines. Each symbol represents a different field of view from two independent days of imaging. n>75 nuclei per condition. (E) Refers to images in (B). Scatter plot shows the ratio of total nuclear fluorescence intensity of Sun2C over GFP for individual cells from different mutant lines and normalized to average ratio of WT. Data pooled from two independent experiments. n> 400 cells per condition. *t-test* was performed to measure significance against WT. (F) Line plot shows relative abundance of SUN2 (normalized to aTubulin) in a Cycloheximide (CHX) time course experiment. Data obtained from densitometry analysis of protein bands on western blots probed with SUN2 (for endogenous SUN2) or GFP (for GFP-SUN2 WT and mutants) antibodies. Each cell line is represented by a differently colored line and data normalized to mean value at 0hr for that cell line. Symbols represent mean values of three independent experiments and bars represent SD. For all scatter plots, red bar shows mean and black bar shows standard deviation. Statistical significance represented as: * p<0.05, ** p<0.01, **** p<0.0001. Values show mean percentage of that group. All images are max intensity projections of confocal z-stacks. All scalebars are 20um.

To identify the cysteines responsible for Sun2C epitope masking, we performed IF with Sun2C antibody in these transgenic lines and found that C719A showed greater binding of Sun2C Ab compared to WT or C577A with significantly higher Sun2C intensity (Fig 3B,3E). Lastly, to precisely map the Sun2C epitope, we expressed a C-terminal 17 AA deletion mutant (1-714AA) and found that Sun2C failed to recognize this construct (Fig 3B). Taken together, these data show that Sun2C Ab recognizes the last 17 amino acids of SUN2 that harbors the C719 residue. Excessive binding of Sun2C Ab to C719A, clearly demonstrates that in *in situ* experiments, Sun2C exclusively binds to SUN2 molecules with reduced cysteines at 719 and 615.

Disulfide bonds in proteins help with the folding and stabilization of specific conformations(Mamathambika and Bardwell, 2008). To address the impact of cysteine mutation on SUN2 protein stability, we compared the protein turnover of SUN2 WT and mutants by treating cells with cycloheximide, which inhibits protein synthesis to determine the residual protein at different time points by immunoblotting. Endogenous and overexpressed SUN2 WT degraded at similar rates and about 50% protein remained after 8 hrs (Fig 3F, black and red lines). Comparatively, both cysteine mutants, C719A and C577A degraded much faster with only 20% and 30% protein remaining after 8 hrs, respectively (Fig 3F, green and blue lines), suggesting that although both cysteines contribute to maintaining SUN2 stability, C615-C719 disulfide bridge is critical for SUN2 protein turnover. Taken together, our data show that the SUN2 terminal disulfide bridge (C615-C719) shields Sun2C Ab epitope that can be revealed upon disrupting this bond (Fig 3G). More importantly, we show that this bond is required for proper localization and turnover of SUN2 thereby providing the first functional evidence for the conservation of Cys615 and Cys719 residues.

### SUN2 cysteines contribute to proper assembly of the LINC complex

Since SUN proteins are an integral part of the LINC complex, we asked whether compromising SUN2 stability upon cysteine mutation, had an impact on the associated LINC complex(Ketema et al., 2007; Stewart-Hutchinson et al., 2008; Lombardi et al., 2011). To test this, we transiently depleted endogenous SUN2 and stained for SYNE3 or EMERIN in WT and mutant expressing lines. In C2C12 MBs not expressing any transgene, SUN2 depletion had little to no effect on SYNE3 or EMERIN (Fig 4A, 4B), suggesting that SUN1 likely compensates for the loss of SUN2 to maintain LINC complexes at the NE. Similarly, overexpression of WT or C577A showed negligible effect. However, as expected, we observed a dramatic loss of both proteins at the NE in cells overexpressing C719A mutant (Fig 4A,4B), showing that C719A acts as a dominant negative to disrupt LINC complexes.

**Figure 4:**
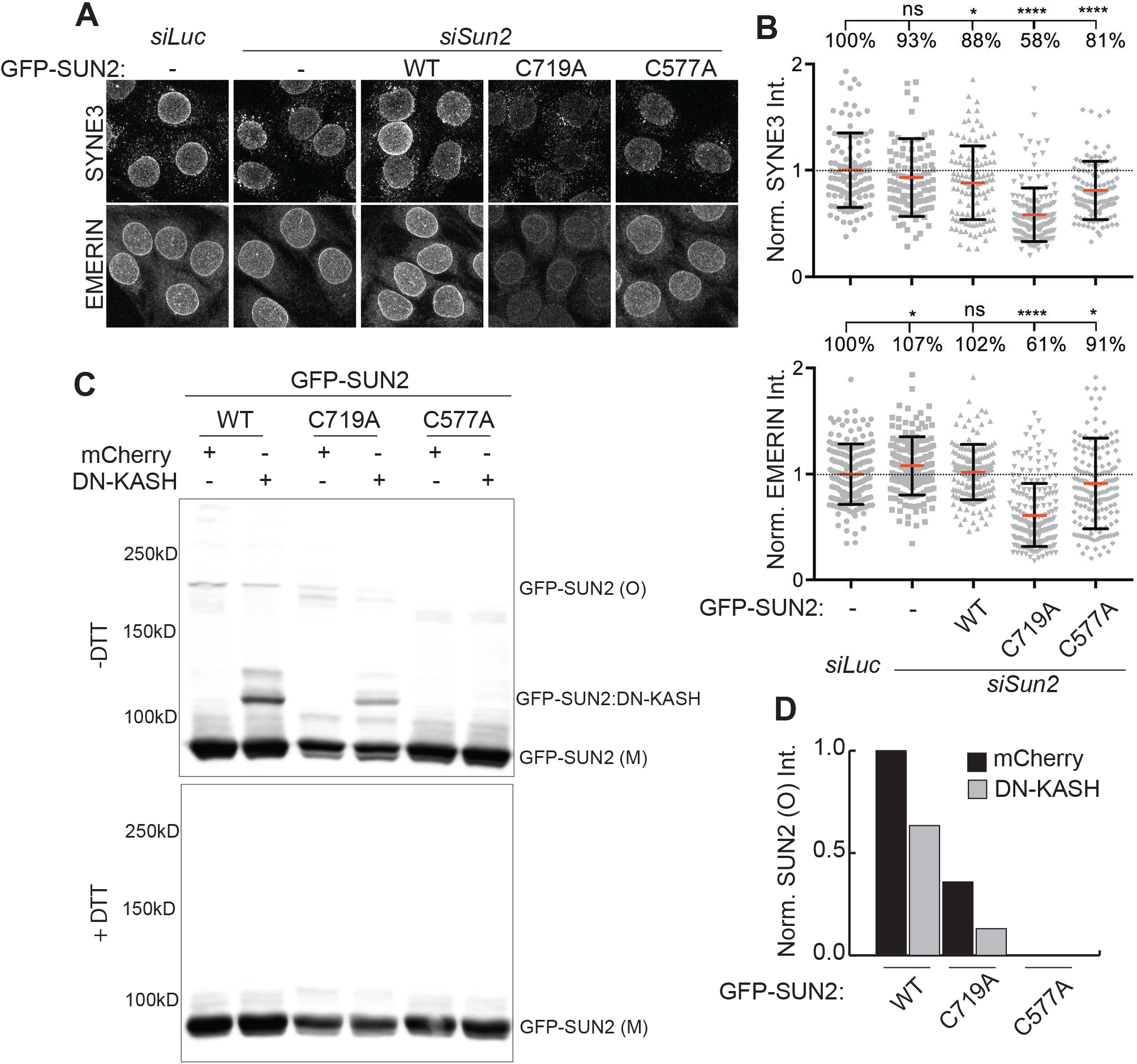
SUN2 Cysteines regulate proper assembly of the LINC complex. (A) Representative confocal images of C2C12 cell lines stably expressing SUN2 WT or mutant constructs, treated with *siLuc* or *siSun2* and stained for either SYNE3 or EMERIN. (B and C) Quantitation for Images in (A). Scatter plot represents total nuclear fluorescence intensity for individual cells across different cell lines. Data pooled from two independent experiments, n>100 cells per condition. One way ANOVA with multiple comparisons was applied to calculate statistical significance against *siLuc*. For scatter plots, red bar shows mean and black bar shows standard deviation. Statistical significance represented as: * p<0.05, ** p<0.01, **** p<0.0001. Values show mean percentage of that group. All images are max intensity projections of confocal z-stacks. Scalebar is 20um. (C) Western blot shows differential migration of ectopically expressed GFP-SUN2 WT and mutant proteins as monomers (GFP-SUN2 (M)) and Oligomers (GFP-SUN2 (O)) in non-reducing (-DTT, top panel) and reducing (+DTT, bottom panel) conditions. GFP-SUN2:DN-KASH complex can be detected as a single band (GFP-SUN2:DN-KASH). (D) Quantitation for (C). Bar graph shows ratio of GFP-SUN2(O) to GFP-SUN2(M) band intensity per condition across SUN2 WT and mutant lines; normalized to WT mCherry expressing cells. Black bars: mCherry and gray bars: DN-KASH expression.

Next, we addressed which of the SUN2 cysteines participate in SUN2 oligomerization and KASH binding. We overexpressed mCherry and DN-KASH constructs in GFP-SUN2 WT and cysteine mutant lines(Sosa et al., 2012) and found that WT showed a single oligomer band and the band intensity decreased upon DN-KASH expression similar to endogenous SUN2 (Fig 4C,4D). Conversely, C719A mutation impaired proper oligomer assembly as evident by the presence of multiple faint bands. Interestingly, C577A mutation completely abrogated SUN2 oligomerization (Fig 4C) suggesting that C577 is critical for the formation of these disulfide-linked oligomer species as well as SUN2:KASH interaction. Overall, our data point towards the importance of these conserved cysteines in proper cellular localization and assembly of LINC complexes specifically at the NE.

### SUN2 cysteines regulate the Actin Cytoskeleton

LINC complexes directly associate with the cytoskeleton to balance mechanical forces across the cell and thereby influence cytoskeletal remodeling(Gimpel et al., 2017; Lombardi et al., 2011; Stewart et al., 2015).

To address the role of SUN proteins in regulating actin cytoskeletal organization, we transiently depleted SUN proteins in C2C12 MBs and visualized polymerized actin (F-actin) (Fig 5A). Surprisingly, we observed that any perturbation to either SUN1 or SUN2 caused a significant decrease in F-actin intensities (~32% loss), which declined even further upon loss of both SUN proteins (Fig 5A,5C). This data suggests that Factin is sensitive to the level of SUN proteins and that neither of the SUN proteins can compensate for maintaining proper F-actin levels. Next, to test whether SUN2 cysteines additionally contribute to cellular F-actin regulation, we transiently depleted endogenous SUN2 in SUN2 WT and cysteine mutant lines and visualized F-actin structures (Fig 5B). Our F-actin quantification revealed that, although endogenous SUN2 depletion had minor but significant effect in WT line; both Cysteine mutants (C719A and C577A) had a dramatic loss of F-actin (~ 50% decrease) with levels comparable to *siSun1/2* double knockdown (Fig 5B, 5D). Taken together, our data show that actin cytoskeleton regulation is not only sensitive to the total level of SUN proteins but also depends on their redox state.

**Figure 5:**
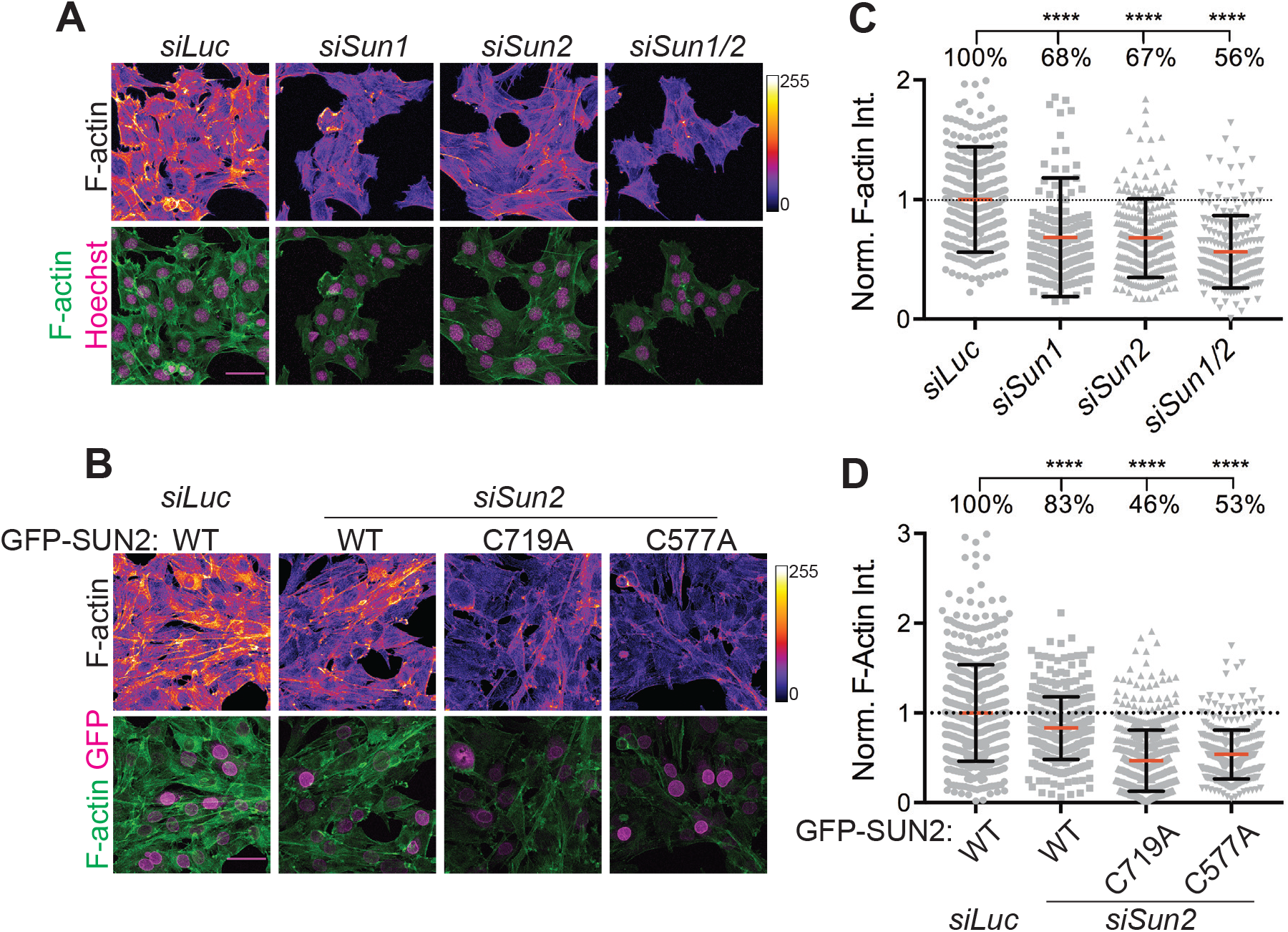
LINC complexes regulate actin cytoskeletal structure. (A) Representative confocal images of C2C12 cells, 72hr post treatment with siRNA against *Luciferase* (control), *Sun1*, *Sun2* and *Sun1/2*, stained with fluorescently labelled Phalloidin for F-actin (pseudocolored, top panel) and nuclei counterstained with Hoechst (bottom panel). (B) Representative confocal images of C2C12 cell lines stably expressing GFP-tagged WT or mutant constructs, subjected to *siLuc* or *siSun2* treatment for 72hr and stained with fluorescently labelled Phalloidin for F-actin (pseudocolored, top panel) and GFP (bottom panel). (C and D) Quantitation for A and B respectively. Scatter plot represents the distribution of Phalloidin intensities per cell in each condition. Data pooled from two independent experiments and normalized to average value of *siLuc* (for C) or WT *siLuc* (for D). n>230 cells per condition. One way ANOVA with multiple comparisons was applied to calculate statistical significance against *siLuc* (for C) and *siLuc* WT (for D). For all scatter plots, red bar shows mean and black bar shows SD. Statistical significance represented as: * p<0.05, ** p<0.01, **** p<0.0001 and ns not significant. Values show mean percentage of that group. All images are max intensity projections of confocal z-stacks. Scale bar is 50 um. F-actin was pseudocolored with FIRE lookup table and intensity scale shown on the right side.

### Dynamic changes in SUN2 cysteine oxidation state regulate Cell Migration

The LINC complex plays an important and well documented role in cell migration during wound healing of a cell monolayer(Gant et al., 2010; Zhu et al., 2017; Chang et al., 2015; Cain et al., 2018; Lombardi et al., 2011). We took advantage of wound healing/scratch assay to investigate the dynamics of SUN2 terminal disulfide bond during directional cell movement. To test for disulfide bond rearrangements, we performed IF with Sun2C and Sun2N Abs at different time points post wound. We observed a distinct loss of Sun2C staining specifically in migrating cells at the edge of the wound compared to confluent monolayers as early as 6 hr post wound (Fig 6A). We quantified these changes and observed a significantly lower Sun2C intensity at the wound edge 8 hrs post scratch compared to 0 hr time point (Fig 6B, 6C Edge). Whereas, Sun2C intensity remained unchanged away from the scratch site (Fig 6B, 6C Center). This experiment shows that migrating cells exclusively present at the wound edge undergo dynamic SUN2 terminal cysteine oxidation suggesting a remodeling of LINC complex.

**Figure 6:**
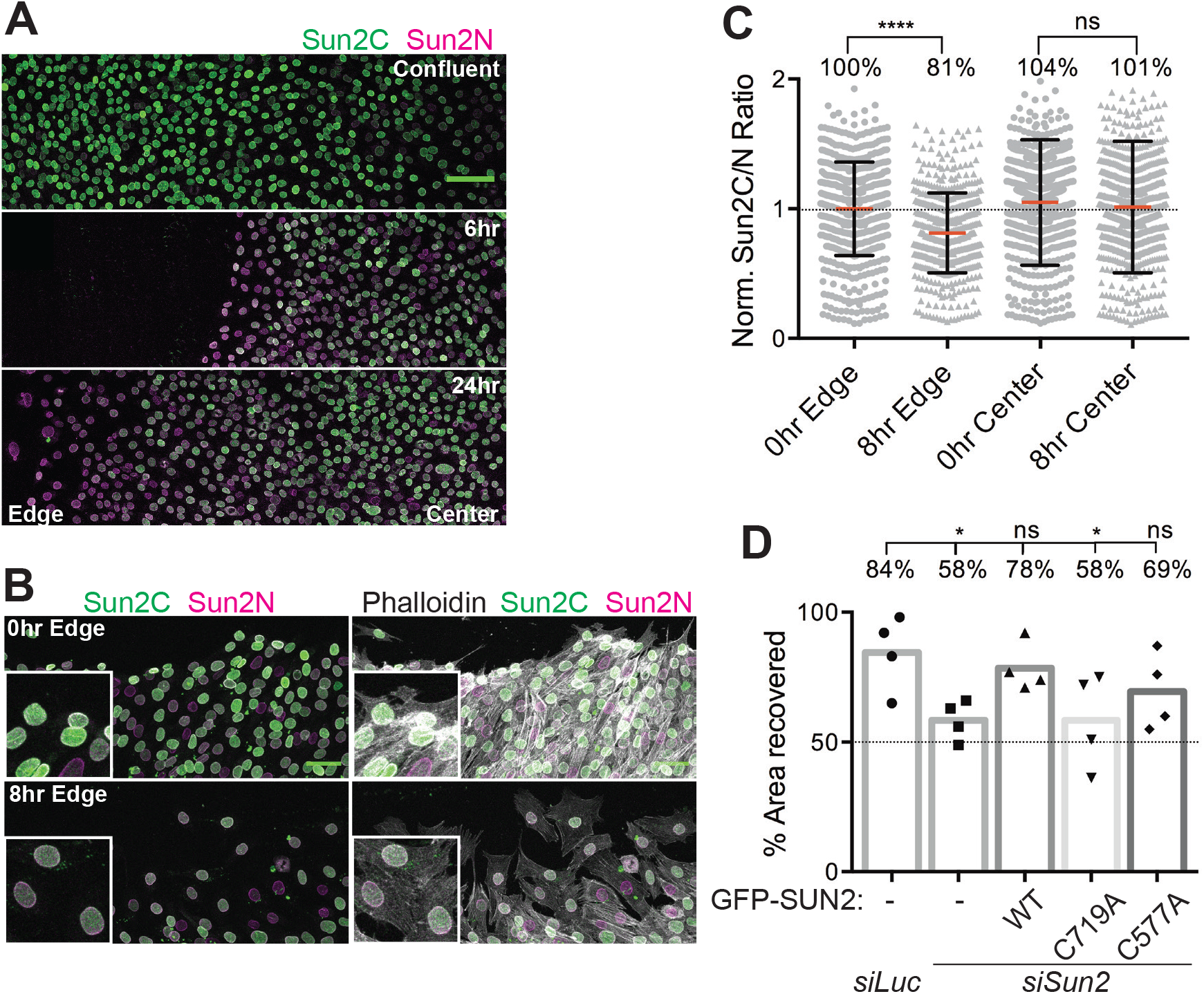
Dynamic regulation of SUN2 terminal disulfide bond during cell migration. (A) Stitched confocal image panels of confluent C2C12 cell monolayers (top panel) in a wound healing assay, 6hr (middle panel) and 24hr (bottom panel) post-wound and co-stained with Sun2C and Sun2N Abs. Wounded and un-wounded regions are labelled as Edge and Center respectively. (B) Representative confocal image of wound healing assay. Images taken 0hr and 8hrs post wound and stained with Sun2C, Sun2N and Phalloidin. Inset shows zoomed section of the image. (C) Quantitation for images in (B). Scatterplot shows the ratio of total nuclear fluorescence intensity of Sun2C over Sun2N for individual cells at 0hr or 8hr post wound, at the edge of the wound (Edge) or unwounded region (Center). Data pooled from two independent experiments where n>500 cells per condition. red bar shows mean and black bar shows standard deviation. Statistical significance tested using *t-test*. (D) Bar graph shows mean percentage of area recovered 24hrs post wound in a wound healing assay for different C2C12 engineered cell lines. Each symbol represents individual replicates of four independent experiments. One way ANOVA was performed with multiple comparisons against *siLuc* sample. Statistical significance represented as: * p<0.05, ** p<0.01, **** p<0.0001, ns not significant. Values show mean percentage of that group. All images are max intensity projections of confocal z-stacks. All scalebars are 100um.

To directly address the importance of SUN2 cysteines during cell migration, we performed wound healing assays in our SUN2 WT and cysteine mutant lines upon transient depletion of endogenous SUN2. Our data shows that cells migrate slower in the absence of SUN2 but recover upon compensating with WT and to a lesser extent with C577A mutant. However, cells overexpressing C719A exhibited significantly slower migration compared to the control *siLuc* (Fig 6D), suggesting that SUN2 terminal disulfide bridge is required for proper cell migration. Consistent with previous reports, our data clearly demonstrates that upon receiving migratory cues, cells reorganize their nucleo-cytoskeletal structure and engage their LINC complexes to facilitate directional cell movement. For the first time, these transition states can be clearly visualized using Sun2C antibody at precise spatio-temporal resolution providing molecular insights into cell migration.

### Stress-related regulators of SUN2 terminal disulfide bond

Our data directly links the redox state of SUN2 terminal cysteines to several molecular and cellular phenotypes. The C-terminus of SUN2 resides in the ER/NE lumen that harbors proteins crucial for protein folding, quality control, intracellular calcium homeostasis, and redox balance(Görlach et al., 2006). Therefore, we tested whether SUN2 cysteines respond to changes in ER homeostasis. To induce ER stress, we transiently treated C2C12 MBs with either an ER Ca^2+^ ATPase inhibitor, Thapsigargin (Tg) that causes loss of calcium from ER lumen, or 16F16 (iPDI), a Protein Disulfide Isomerase (PDI) enzyme inhibitor that compromises protein folding in ER(Hoffstrom et al., 2010). Upon IF with Sun2C and Sun2N Abs, we observed a significant increase in Sun2C signal in Tg (Fig 7A, 7B) and iPDI (Fig 7C) treated cells compared to DMSO control. These changes were observed within 1 hour of treatment and did not change with increase in treatment time suggesting that SUN2 terminal cysteines are highly sensitive to ER environment and quickly undergo reduction upon perturbing ER homeostasis.

**Figure 7:**
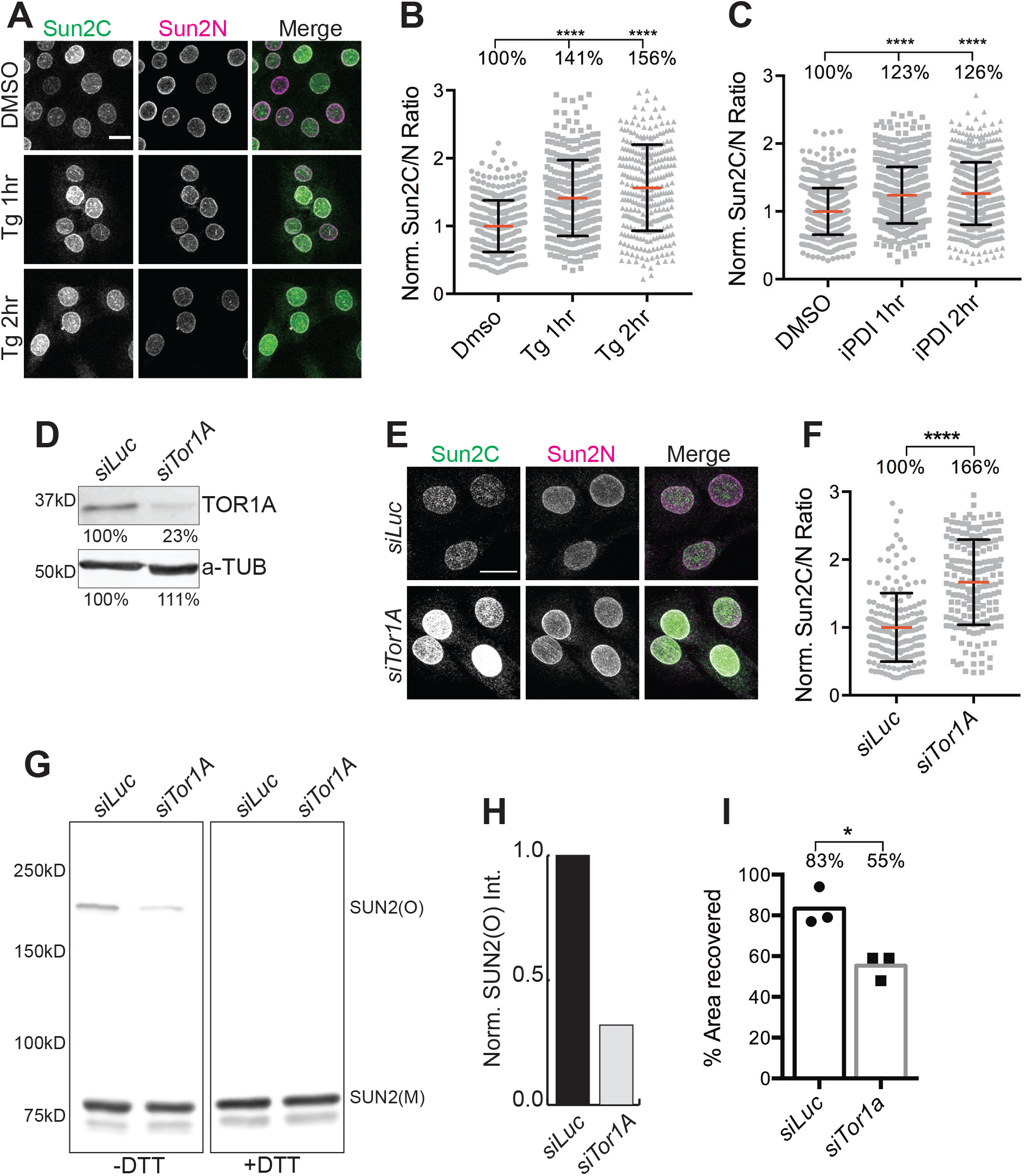
Endoplasmic reticulum environment regulates oxidative state of SUN2 terminal disulfide bond. (A) Representative confocal images of C2C12 MBs treated either with DMSO or Thapsigargin (Tg) for 1-2hrs and co-stained with Sun2C and Sun2N antibodies. (B and C) Scatter plot shows the ratio of total nuclear fluorescence intensity of Sun2C over Sun2N for individual cells in DMSO control, Tg treated (for B) and Protein Disulfide Isomerase inhibitor treated (iPDI, for C); normalized to average ratio of DMSO. Data pooled from two independent experiments. n>250 (for B) and n>600 (for C) cells per condition. (D) Western blot against Torsin1A (TOR1A) and a-Tubulin (a-TUB) in Luciferase control (*siLuc*) or *Tor1A* (*siTor1A*) treated MBs. Values show relative band intensity. (E) Representative images of *siLuc* or *siTor1A* treated MBs co-stained with Sun2C and Sun2N antibodies. (F) Quantitation for images in (E). Scatter plot shows Sun2C/N ratio in *siLuc* and *siTor1A* treated cells normalized to average ratio of *siLuc*. Data pooled from two independent experiments. n> 180 cells counted per condition. (G) Western blot shows SUN2 oligomers (SUN2(O)) and monomers (SUN2(M)) in *siLuc* and *siTor1A* treated cells in non-reducing (-DTT) and reducing conditions(+DTT). (H) Quantitation for (G). (I) Bar graph shows mean percentage of area recovered 24hrs post wound in a wound healing assay for C2C12 cells treated with either *siLuc* or *siTor1A*. Each symbol represents individual replicates of three independent experiments. For all scatter plots, red bar shows mean and black bar shows SD. *t-test* was applied to calculate statistical significance against controls (DMSO for (B) and (C); *siLuc* for (F) and (I)). * p<0.05, ** p<0.01, **** p<0.0001. Values show mean percentage of that group. All images are max intensity projections of confocal z-stacks. All scalebars are 20um.

Multiple studies have implicated Torsin1A (TOR1A) in remodeling LINC complexes at NE(Nery et al., 2008; Saunders et al., 2017), however, the molecular mechanisms remain unclear. TOR1A is part of AAA+ ATPase superfamily of protein that specifically resides in ER/NE lumen and acts as a molecular chaperone to assemble protein complexes(Burdette et al., 2010; Laudermilch and Schlieker, 2016). Therefore, we asked whether TOR1A participates in maintaining the redox state of SUN2 terminal cysteines. We transiently depleted TOR1A in C2C12 MBs (Fig 7D) and upon co-staining with both SUN2 antibodies, we observed a dramatic increase in Sun2C signal in TOR1A depleted cells (Fig 7E, 7F). This suggests that loss of TOR1A leads to reduction of SUN2 terminal cysteine residues. Moreover, we observed a dramatic loss of C577 linked SUN2 oligomers in TOR1A depleted cells (Fig 7G, 7H). To evaluate the functional significance of these observations, we performed a wound healing assay and found that TOR1A depleted cells migrated slower and showed a significant decline in wound closure compared to control (Fig 7I). This data is consistent with a previous study (Nery et al., 2008) and supports our own observations showing relatively slower migration of C719A lines lacking SUN2 terminal disulfide bond. Taken together, our data show that the redox state of terminal cysteines in SUN2 could dynamically be altered, not only by TOR1A, but by additional factors that regulate ER homeostasis. These results have major implications in further understanding the rapid remodeling of LINC complexes at the NE.

## Discussion

### Conformation-specific SUN2 antibody

Conformation-specific antibodies are indispensable tools for *in situ* biological research as they provide unprecedented insights into the function of proteins in their physiological environment. Structural proteins like Integrins and Lamins are excellent examples where conformational epitopes have unraveled structure-function relationships(Dyer et al., 1997; Ihalainen et al., 2015; Bazzoni et al., 1995). Here, we characterize a SUN2 Ab (Sun2C) that has been widely used in literature and show, for the first time, that it is conformation-specific. At the molecular level, we find that SUN2:KASH interaction leads to C615-C719 disulfide bond formation in SUN2, that masks Sun2C epitope. Using a quantitative imaging assay, we demonstrate that Sun2C epitope masking occurs during key cellular processes like cell proliferation, differentiation and migration. Since LINC complexes have been previously implicated in all these processes(Aureille et al., 2019; Carley et al., 2021; Gant et al., 2010; Chang et al., 2015; Déjardin et al., 2020), our data complements these studies by providing a new marker for SUN2:KASH interaction *in situ* and has the potential to generate novel insights from new and existing datasets. Additionally, our findings support the idea that instead of changing total protein levels at the NE; LINC complexes can rapidly contribute to biological function by dynamically altering their conformation.

### SUN2 disulfide bonds and LINC complex formation

*In vitro* structure data has shown that SUN2 terminal disulfide bond remains intact in KASH bound and unbound state(Sosa et al., 2012; Wang et al., 2012; Cruz et al., 2020). Our data in MBs reveals that newly synthesized SUN2 in the ER and KASH bound SUN2 at the NE do not show any Sun2C staining, providing *in vivo* confirmation of previous structural data. Interestingly, we observe Sun2C staining at the NE in proliferating MBs that suggests that terminal disulfide bond is dynamic and alters redox state at the nuclear periphery. Our functional experiments show that Sun2C staining is enhanced when SUN2 is unbound and devoid of KASH suggesting that the terminal disulfide bond is responsive to the SUN2:KASH interaction and a fraction of SUN2 exists naturally in a KASH unbound state. The presence of reduced SUN2 cysteines specifically at the NE is intriguing and highlights the importance of C615-C719 disulfide bond. We show that disrupting a single disulfide bond by mutating C719 results in accumulation of the protein in the ER, enhanced rate of degradation, and disruption of LINC complex assembly and function. Moreover, we find improperly assembled SUN2 C719A homo-oligomers on non-reducing gels. This suggests that terminal disulfide bond is required for proper protein folding in the ER, regulation of protein-protein interactions, and participation in proper INM localization of SUN2. So why are SUN2 cysteines reduced at the NE? Our Nesprin-3 KD and SUN1L OE experiments show that the lack of KASH engagement is sufficient to reduce terminal disulfide bond in SUN2, suggesting the presence of an active disulfide rearrangement mechanism at the NE.

At the mechanistic level, it has been shown that SUN2 C577 forms a disulfide bond with KASH C-23 to strengthen SUN-KASH interactions(Sosa et al., 2012; Cain et al., 2018; Jahed et al., 2015). We show the existence of disulfide linked SUN2 homo-oligomers that disappear upon KASH interaction on non-reducing gels. Using a mutagenesis approach, we identify C577 as the critical residue for SUN2 homooligomerization. Since the same residue is involved in homo-oligomerization and KASH interaction, it is imperative that SUN2 oligomers undergo disulfide bond rearrangement to switch partners. However, structural studies show C577 independent SUN2 oligomerization and KASH engagement(Cain et al., 2018). Therefore, the exact nature of disulfide linked SUN2 homo-oligomers remains to be further characterized. Considering that only a small fraction of total SUN2 exists as a disulfide linked oligomer, we think these might be either very short-lived intermediates or remnant of higher order branched LINC assemblies connected through C577 disulfide bonds(Gurusaran and Davies, 2021). In addition, coiled coil domains in SUN2 contribute to oligomerization *in vitro*(Nie et al., 2016), which cannot be resolved by non-reducing gels, and require future investigation. Consistent with previous studies, we find that mutating C577 does not affect protein localization, but increases protein degradation that results in significant effect on LINC complex assembly and function. Most importantly, C577A mutant shows an increase in Sun2C signal suggesting that loss of interaction between SUN2 and KASH cysteines, does affect the SUN2 terminal disulfide bond. Similarly, C719A mutant shows misassembled C577 linked homo-oligomers. This shows a dynamic interplay among all three highly conserved cysteines in SUN2 and supports the idea that SUN2 undergoes a disulfide bond rearrangement at the NE during LINC complex formation. Based on these observations, we propose a revised model of LINC assembly and disassembly from the point of view of disulfide bond rearrangement (Figure 8). Overall, our analysis reconciles *in vitro* data with *in vivo* observations and strives to fill a major gap in our understanding of LINC complex structure and function *in vivo*.

**Figure 8:**
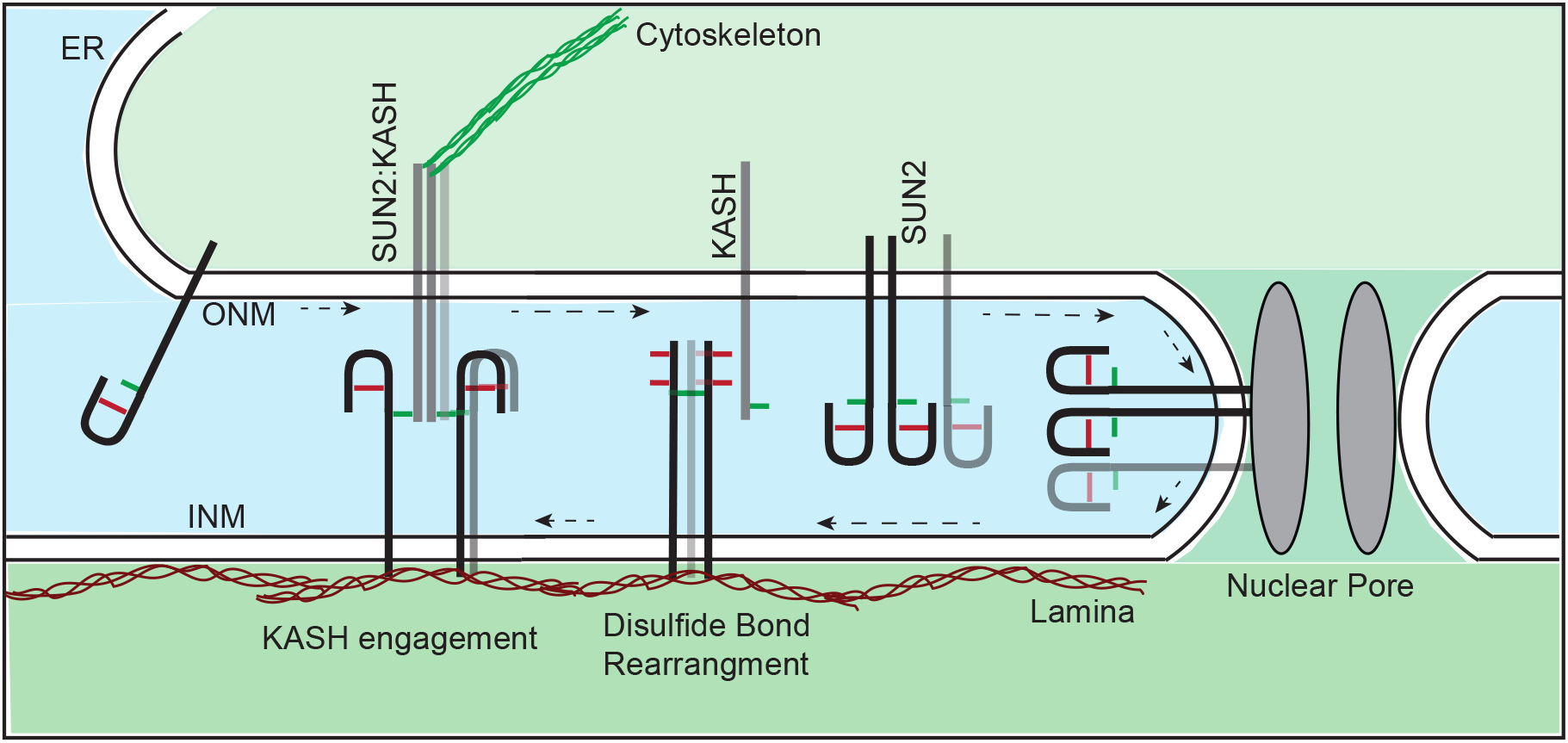
A proposed model for the LINC complex assembly at the NE. Model shows the redox state of SUN2 (black/gray bar) cysteines C719-C615 (red) compared to C577 (green) during transition from ER/ONM to INM (black arrows) and subsequent assembly of SUN2:KASH LINC complex at the INM.

### SUN2 disulfide bonds in cytoskeleton organization and cell migration

LINC complexes interact with the cytoskeleton and affect its organization. Recent literature implicates SUN1 and SUN2 in actin cytoskeletal regulation(Gant et al., 2010; Porter et al., 2020; Stewart et al., 2019). However, whether this regulation is directly mediated through LINC complex or indirectly through signaling pathways remains unclear. Our data show that loss of either of the SUN proteins decreases polymerized actin levels suggesting that SUN1 and SUN2 do not completely compensate each other to maintain F-actin levels in C2C12 MBs. Importantly, we find that both cysteines (C719 and C577) contribute to actin cytoskeleton maintenance as mutating these cysteines leads to dramatic loss in F-actin levels. For C719A, this phenotype can be explained by a concurrent loss in other LINC member (SYNE/EMD). However, Factin loss in C577A was surprising since we did not see much change in Nesprins. Interestingly, according to recent reports, SUN2 seem to regulate RhoA signaling pathway that controls actin polymerization and mechanosignaling, but the mechanisms remain unclear(Thakar et al., 2017; Mu et al., 2020; Porter et al., 2020). This raises a possibility of SUN2 regulating actin independent of LINC complex. Another possibility is that C577A does not affect total levels of Nesprin but decreases the residence time of Nesprins on the NE thereby compromising the stability of Nesprin-Actin interaction(Saunders et al., 2017). Indeed, our data show that C577A degrades faster compared to WT. Taken together, our study points towards the importance of SUN2 Cysteine residues and their redox states in the maintenance of proper actin cytoskeleton system, although further experiments are required to understand the mechanisms. Since Factin is directly related to mechanical stiffness and fluidity of the cell, these findings have implications in cellular transformation, gain of invasive properties and diagnostic potential in disease conditions(Kalukula et al., 2022).

Directional migration of cell monolayer has been well documented and requires SUN2 dependent LINC complex and actin cytoskeleton(Gant et al., 2010; Zhu et al., 2017). Our 2D migration assay shows that SUN2 terminal disulfide bond is intact in cells migrating at the wound edge and disrupting this bond negatively affects migration but does not completely abolish it. This data is consistent with Live FRET measurements of migrating cells that show Nesprins under increased tension at the leading edge of the wound(Déjardin et al., 2020). Altogether, our work provides visual molecular insights into *in vivo* LINC complex dynamics during cell migration. We propose, in future, Sun2C antibody could be used to investigate SUN-KASH engagement during neuronal and muscle development, cancer cell migration, mechanobiology, and other processes where LINC complexes have already been shown to play an important role.

### SUN2 Disulfide regulation

Principles behind proper assembly and disassembly of LINC complexes at the NE remain elusive. We show that SUN2:KASH interaction leads to SUN2 terminal disulfide formation and the loss of this bond accelerates SUN2 protein turnover. Structure and molecular simulation dynamics studies have pointed to the importance of pH and ion concentration for *in vitro* LINC assembly(Jahed et al., 2018; Gurusaran and Davies, 2021). We show that perturbing ER homeostasis is sufficient to rapidly reduce SUN2 terminal cysteines *in* situ thereby demonstrating that ER microenvironment chemically modifies SUN2. Post-translational modifications of SUN2 have been known to impact its function(Gilbert et al., 2019), therefore, our data suggest an intriguing possibility of a redox dependent remodeling of LINC complexes. Several studies have pointed to Torsins as regulators of LINC complexes. TOR1A interacts with SUN and NESPRIN proteins(Chalfant et al., 2019; Nery et al., 2008) and seems to regulate their NE positioning and mobility, however, the mechanisms are not clearly understood. Our data shows that the depletion of TOR1A leads to the loss of SUN2 terminal disulfide bond, SUN2 homo-oligomers and affects cell migration. Since TOR1A also participates in regulating ER stress response(Chen et al., 2010), it is unclear whether TOR1A directly or indirectly contributes to SUN2 cysteine oxidation. Overall, this study lays the foundation for future identification of redox-based LINC complex regulators. Moreover, this work supports the idea that, in addition to their roles in protein folding and stability, some disulfide bridges may possess key regulatory property that diversifies their function(Hogg, 2003).

## Materials and methods

### Cell culture and cell line generation

Mouse C2C12 myoblast cells were cultured in Growth media (DMEM with 20% FBS and 1% Pen-Strep) and passaged every other day. For differentiation to myotubes, confluent C2C12 myoblasts were cultured in Differentiation media (DMEM with 2% Horse serum and 1%Pen-Strep) with media changes every other day. Cell lines cultured under 37°C and 5%CO2 incubator conditions. C2C12 cells stably expressing GFP-SUN2 WT, C719A and C577A constructs were generated by retroviral transduction of C2C12 cells. For retroviral packaging and production, 293T cells were transfected with 4ug of Ampho, 4ug of plasmid of interest and 32ul of Polyethylenimine (PEI) in 10cm plates. Virus containing media was harvested at 48hrs and cell free supernatant was used to transduce early passage C2C12 cells. Successfully transduced cells were selected by culturing in media containing 10ug/ml Blasticidin for 4 days. Selected cells were expanded in growth media and multiple vials were frozen for future use.

### Plasmids and siRNA

N-terminal GFP-tagged SUN2 wild type (WT) expression plasmid was constructed as previously described (Buchwalter et al., 2019) using a mouse SUN2 sequence (Uniport ID Q8BJS4). GFP and SUN2 open reading frames were individually amplified using primers with overlapping regions between GFP and SUN2. Forward primer for GFP and reverse primer for SUN2 included attB sites for Gateway cloning. Next, using an overlap PCR strategy GFP and Sun2 amplicons were stitched together {Heckman and Pease, 2007} to generate a single amplicon with attB sites. Using a Gateway cloning kit, the GFP-SUN2 amplicon was first cloned into a PDONR207 vector and later into PQCXIB (Campeau et al., 2009), a retroviral backbone gateway destination vector. GFP-SUN2 WT construct was used as a template for cloning all other SUN2 mutant versions. GFP-SUN2(1-714AA) was amplified using GFP forward primer and a reverse primer containing a stop codon after amino acid 714 (5’-ctaGTGGCCCCAGTTGGTCAGGATCC-3’) followed by Gateway cloning into PQCXIB vector. We used Quickchange mutagenesis strategy to generate SUN2 cysteine mutants C719A and C577A. We used an overlapping primer set to amplify GFP-SUN2 WT construct by replacing either TGT(C719) or TGC(C577) to GCC (C719A or C577A) thereby substituting Alanine for Cysteines. All constructs were verified by sequencing. pmCherry and pmCherry -C1:KASH1 (DNKASH) constructs were previously published (Hatch and Hetzer, 2016). A previously published plasmid, pCDH-CMV-MCS-EF1 copGFP-T2A-puro:SS-HA-Sun1L-KDEL (Lombardi et al., 2011)was used to subclone the SS-HA-SUN1L-KDEL region into a pC1 vector to generate pC1:SS-HA-Sun1L-KDEL-IRES-mCherry (SUN1L) construct (plasmid generated by Emily Hatch). pCDH-CMV-MCS-EF1-SS-GFP-KDEL plasmid was a gift from J. Lammerding (The Weill Institute for Cell and Molecular Biology, Cornell Univeristy, Ithaca, NY (Lombardi et al., 2011)

Control siRNA against Luciferase (siLuc) was custom ordered from Life technologies (sense: 5’-uaugcaguugcucuccagcdtdt-3’). Individual or pooled siRNAs against other targets were ordered from Horizon Discovery (Dharmacon), Lafayette, CO. with following catalogue numbers: Sun1 (ON-TARGETplus Mouse Sun1 (77053) siRNA - Individual, J-040715-10); Sun2 (ON-TARGETplus Mouse Sun2 (223697) siRNA – Individual, J-041247-09); Syne3 (ON-TARGETplus Mouse Syne3 (212073) siRNA - SMARTpool, L-052180-01); Tor1a (ON-TARGETplus Mouse Tor1a (30931) siRNA – SMARTpool, L-051579-01).

### Transfections and pharmacological treatments

DNA transfection was performed using Lipofectamine 3000 following manufacturers protocol directly on cells plated on previous day, in ibidi chambers (for Imaging) or 6 well plates (for Biochemistry). Experiments were performed 24hrs post DNA transfection. siRNA (50nM final conc) was reverse transfected in 6 well plates using Lipofectamine RNAiMAX at 0hr and repeated at 24hr. Cells were split at 48hr and either seeded in ibidi chambers (for imaging) or passaged into 6 well plates (for protein) to be analyzed at 72hr time point.

Cells were seeded a day before in ibidi chambers and treated with either DMSO, 100nM Thapsigargin (Life Technologies # T7458) or 50uM iPDI (PDI inhibitor 16F16, Sigma# SML0021) in growth media for 1 or 2hr before fixation and IF. For DTT rescue experiment, cells in ibidi chambers were treated with media (Growth for MBs/ Differentiation for MTs) containing either DI water (Vehicle) or 5uM final conc. of DTT (Acros Organics#AC16568) for 3 minutes before fixation and IF.

For cycloheximide chase experiment, cells were seeded in 6-well plates the previous day and treated either with 200ug/ml Cycloheximide (CHX) (Sigma-Aldrich # C-7698) in fresh growth media for 2, 4 or 8 hrs; or without CHX (0hr). Cells were harvested by trypsinization and flash frozen cell pellets were stored in −20C for future protein analysis.

### Immunofluorescence

Cells were cultured in Ibidi μ-Slide 8 Well (Ibidi #80826) chambers with indicated treatments. On the day of the experiment, cells were washed once with PBS, fixed in 4% Paraformaldehyde in PBS for 5 min, followed by two PBS washes for 5 min. At this point, cells were either stored in fresh PBS at 4C in humidified chamber for future staining or processed further in the following manner. Cells were permeabilized and blocked in IF Buffer (10mg/ml BSA, 0.1% Triton X-100, 0.02% SDS in PBS) for 20 min followed by incubation in IF buffer containing primary antibodies at appropriate dilution for two hours at RT. Cells were washed three times in IF buffer followed by incubation in IF buffer containing secondary antibodies (and Phalloidin-647 when appropriate) for 1hr at RT. For DNA counterstain, cells were incubated in IF buffer containing Hoechst for 5 min, followed by three washes with IF Buffer. Finally, cells were washed once with PBS and stored in PBS for confocal imaging the same day.

Following primary antibodies and their respective dilutions were used in IF experiments: mouse Sun1 clone 12.10F (MilliporeSigma #MABT892; 1:250); rabbit Sun2(Sun2C) EPR6557 (Abcam #ab124916; 1:500); mouse Sun2(Sun2N) clone 3.1E (MilliporeSigma # MABT880,1:500); mouse Nesprin1 MANNES1A(7A12) (ThermoFisher #MA5-18077, 1:500); mouse Nesprin3 (Nordic-MUBio # MUB1317P,1:250); rabbit Emerin D3B9G XP (CST # 30853); mouse MHC MF-20 (in house, Developmental Studies Hybridoma Bank at the University of Iowa). For Actin staining: Phalloidin Alexa Fluor-647(ThermoFisher #A22287; 1:1000) was used. All Alexa Fluor dye conjugated secondary antibodies against rabbit and mouse were obtained from ThermoFisher and used at 1:2000 dilution.

### Microscopy, image analysis and quantitation

All samples were imaged on Leica SP8 scanning confocal microscope using 63X/1.4 NA or 20x/0.75 NA oil immersion objectives. All samples in a given experiment were imaged using same laser intensities and gain settings without saturation. Multiple z-slices were acquired to image entire nuclei and quantitation was performed on maximum intensity projections of z-stacks in Fiji (ImageJ)(Schindelin et al., 2012). Nuclear fluorescence intensity measurements were performed in Fiji by generating a nuclear mask based on Hoechst DNA staining and applying the mask to other background subtracted channels of the same image. This allowed us to measure the integrated densities for each detected nuclei across different channels in a multicolor imaging experiment. F-actin intensity measurements were performed in CellProfiler (McQuin et al., 2018) by determining primary object as Hoechst-stained nuclei and secondary object as Phalloidin-stained F-actin in order to segment out cell boundaries in each image. Integrated densities were measured in Phalloidin channel for each segmented cell.

For Sun2C/N ratio quantification, Sun2C fluorescence intensities measured using the above method were divided by the corresponding Sun2N intensities per nucleus per condition generating a ‘Sun2C/N ratio’. Ratios from different replicates were pooled together. For normalization across different conditions in a given experiment, all values were divided by the average value of the control group in that given experiment, thereby generating a ‘Norm Sun2C/N’. ‘Norm Sun2C/GFP’ ratios were generated in the same way by using GFP intensity in place of Sun2N. To quantify ‘Norm Int.’ for NE markers in different experiments, we used the Integrated densities per nuclei per condition for that particular NE marker across replicates and normalized it to the average value of the control group in that given experiment. To quantify percentage of cells with SUN2 signal in ER, we took images of cells either stained for endogenous SUN2 or expressing GFP tagged constructs (SUN2 WT and mutants) in 2-3 different field of views on two independent days. We manually counted SUN2/GFP positive cells and determined whether SUN2/GFP signal was restricted to the nucleus or also detected in the ER for that particular cell. Percentage of ER positive signal cells was calculated per field of view and represented as one data point. All calculations were done in MS Excel.

For wound healing assays, brightfield images of 2-3 different wound regions per condition were acquired on Invitrogen EVOS FL system using 4x air objective. Each image was analyzed using Wound_healing_size_tool plugin(Suarez-Arnedo et al., 2020) in Fiji by adjusting the parameters to quantify the percentage cell free region.

Densitometry analysis of western blots was performed using the Analyze Gels function in Fiji. For oligomer gels, Sun2 oligomer (Sun2(O)) band intensity was divided by Sun2 monomer band intensity (Sun2(M)) for that particular sample/lane. Additionally, all other samples were adjusted to control group/ first lane). For Norm. GFP-SUN2 level quantitation, GFP band intensity was normalized to aTUB levels of the same sample. Additionally, all samples were normalized to the average value of GFP-SUN2 WT. To calculate Sun2 relative abundance after CHX treatment, SUN2 (for WT) or GFP (GFP tagged SUN2 variants) band intensity was normalized to aTUB band intensity of that sample and then normalized to 0hr time point of that group/ cell line. For Norm. GFP-SUN2 level quantitation, GFP band intensity from untreated sample (0hr) CHX experiment was used and normalized to aTUB levels of the same sample.

All calculations were done in MS Excel. Prism GraphPad was used to perform statistical analysis and generate graphs. Figures were made in Adobe Illustrator. Schematic illustrations were made either in Adobe illustrator or created with BioRender.com

### Wound healing assay

C2C12 cells were reverse transfected twice with siRNAs on Day0 and Day1 as described above. Cells were trypsinized on Day1 evening and seeded in either one well of a 24 well plate (for live imaging) or ibidi chambers (fixed cell imaging) at a desirable density to obtain a confluent monolayer on the following day. On Day2, cell monolayers were serum deprived by changing media to DMEM containing 2% FBS and P/S. On Day3, starved cell monolayers were scratched using a sterile 1ml micropipette tip, washed twice and maintained in low serum condition. Wounded monolayers were imaged on Day3 post-wound for 0hr time point and Day4 for 24hr time point to assess cell free region. For IF staining in wounded monolayers, multiple ibidi chambers were prepare and fixed at 0hr, 6hr or 8hr post wound.

### Protein extraction and western blotting

C2C12 cells expressing different SUN2 mutants were transfected either with mCherry control or DN-KASH expressing plasmids for 24hr and total cell lysates were resolved on reducing/non-reducing gels and western blots probed with GFP antibody to detect the mutant proteins.

For resolving oligomers on non-reducing gels, cell pellets were resuspended in 1x low SDS RIPA buffer (25mM Tris HCl pH 7.6, 150mM NaCl, 1% NP40 substitute, 1% Sodium Deoxycholate, 0.1% SDS) supplemented with 1x cOmplete Protease inhibitor cocktail, 20mM N-Ethylmaleimide (Sigma-Aldrich #E3876) and 250U/ml Benzonase nuclease (Sigma# E8263) and kept on ice for 15 min. Lysates were gently passed through a 27 gauge syringe for 10 times and left on ice for another 15 min. Cell lysates were cleared by centrifugation at 14,000 rpm for 15 min at 4C. The obtained supernatant was divided into two equal parts and either combined with an equal volume of 2x Sample buffer (125mM Tris HCl pH 6.8, 10% Glycerol, 0.01% Bromophenol Blue and 0.1% SDS) or with 2x Sample buffer supplemented with 100mM DTT. +/-DTT samples were then resolved on 6% SDS-PAGE without boiling the samples. Adapted from (Lu et al., 2008)

All other protein extracts were prepared by resuspending cell pellets in Harsh lysis buffer (10mM Tris HCl pH 7.5, 150mM NaCl, 0.2% NP-40, 0.25% sodium deoxycholate, 0.05% SDS, 1mm EDTA) supplemented with 1x cOmplete Protease inhibitor cocktail and 250U/ml Benzonase nuclease(Franks et al., 2016). Cell lysates were passed through 27 gauge syringe for 10 times on ice and lysates were clarified by spinning at 14,000 rpm for 15 min at 4C. Supernatant was transferred to a new tube, combined with SDS loading buffer and heated at 95C for 10 min. Equal amount of cell lysates were resolved on 10% SDS-PAGE.

For western blotting, proteins were transferred to a nitrocellulose membrane using a wet transfer. Membranes were blocked in TBST (0.25% Tween 20, 20 mM Tris, pH 8.0, and 137 mM NaCl) containing 5% non-fat milk for 30 min followed by incubation with primary antibodies overnight in a shaker at 4C. Membranes were washed in TBST thrice; incubated with secondary antibodies for 1 hr at RT followed by three washes with TBST and stored in PBS. Membranes were developed using ECL and signal detected using KwikQuant Imager (KindleBiosciences LLC) or Odyssey Imaging System (LI-COR Biosciences).

Following antibodies and corresponding dilutions were used: mouse anti-Sun2 (Sun2N) clone 3.1E (MilliporeSigma #MABT880,1:1000); rabbit anti-GFP (EnquireBio#ab290-50ul, 1:1000); rabbit anti-Torsin A/DYT1 (TOR1A) (Abcam #ab34540, 1:500); mouse anti-alpha-Tubulin (aTUB) (Sigma-Aldrich #T5168, 1:4000). HRP conjugated secondary antibodies against mouse proteins were obtained from ThermoFisher and used at 1:5000 dilution.

## Supporting information

Sharma Hetzer 2022_Supplementary data

## Acknowledgments

The authors would like to thank Prof. Daniel A. Starr (UC Davis) for helpful discussions, Jan Lammerding’s lab (Cornell) for sharing plasmids and members of the Hetzer lab for helpful comments on the manuscript. This work was supported by the NOMIS foundation (M.W.H), and the National Institutes of Health (R01 NS096786 to M.W.H.).

The authors declare no conflicting financial interests.

---

**Supplementary Figure 1: Antibody validation and knockdown efficiency data.** (A) Representative images of C2C12 cells 72 hr post transfection of siRNA against *siLuc* and *siSun2* co-stained with Sun2C and Sun2N antibodies. (B and C) Quantitation for images in (A). Scatter plot shows total nuclear fluorescence intensities for Sun2C (B) and Sun2N (C) in *siLuc* and *siSun2* treated cells, normalized to average *siLuc* intensity. n> 100 cells per condition. (D) Representative images of C2C12 cells 72 hr post transfection of siRNA against *siLuc, siSun1* or *siSyne3* and co-stained with SUN1 and SYNE3 antibodies. (E and F) Quantitation for images in (D). Scatter plot shows total nuclear fluorescence intensities for SUN1 (E) and SYNE3 (F) in different knockdown conditions, normalized to average *siLuc* intensity. n>900 cells per knockdown. For all scatter plots, red bar shows mean and black bar shows standard deviation. *t-test* was applied to calculate statistical significance against controls *siLuc*. * p<0.05, ** p<0.01, **** p<0.0001. Values show mean percentage of that group. All images are max intensity projections of confocal z-stacks. All scalebars are 50um.

**Supplementary Figure 2: Multiple sequence alignment of SUN domain across different species.** Conserved cysteines are marked in color. Cysteines involved in intra-molecular bond formation are marked in red and cysteine involved in KASH interaction and inter-molecular SUN:KASH disulfide bond are marked in green. Mouse SUN2 sequence is highlighted in gray. Numbers represent the amino acid position in the peptide.

