## Supplementary material for "C-terminal disulfide bond in SUN2 regulate dynamic remodeling of LINC complexes at the nuclear envelope": Sharma Hetzer 2022_Supplementary data

Supplementary Figure 1: Antibody validation and knockdown efficiency data

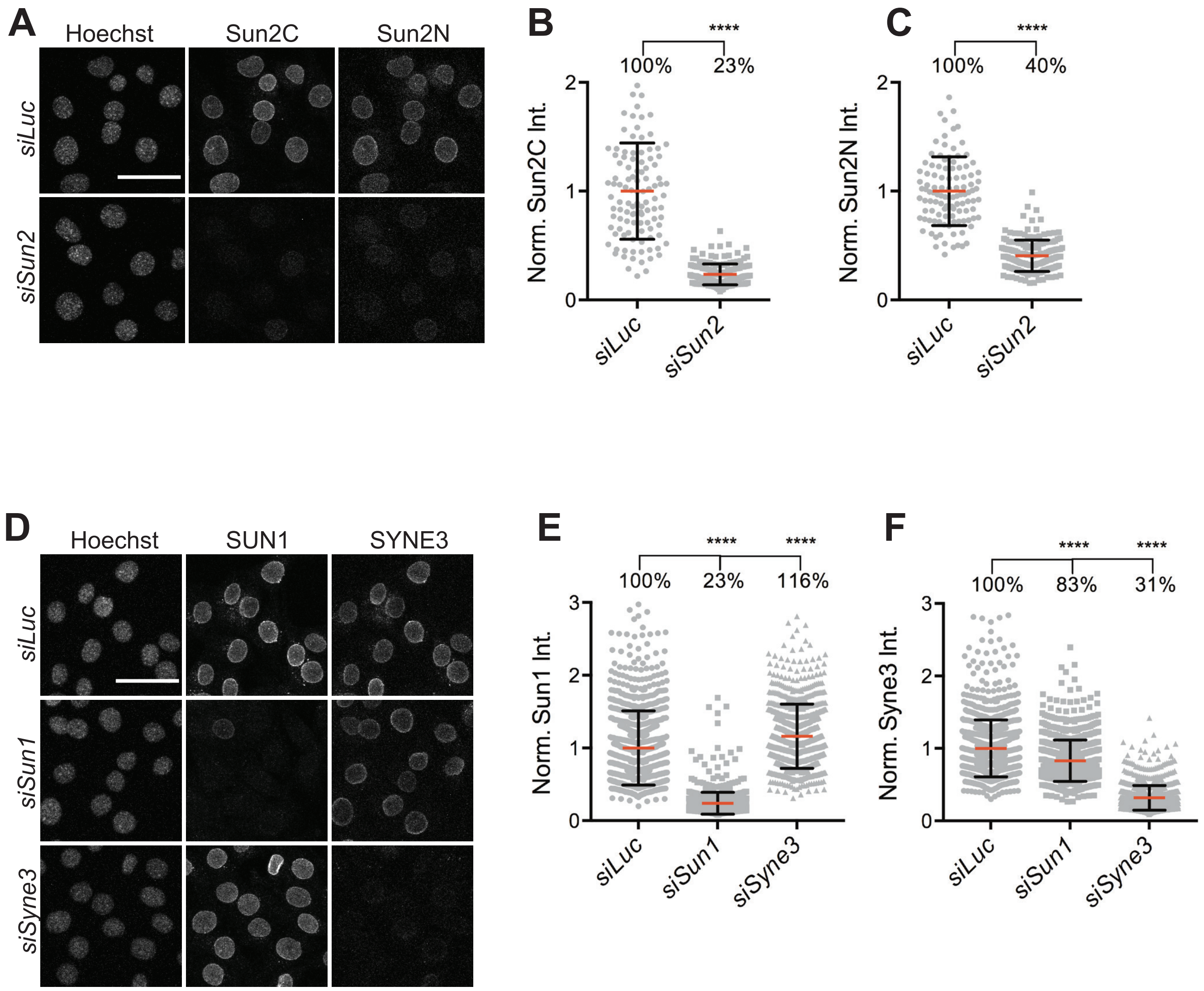

Supplementary Figure 2: Multiple sequence alignment of SUN domain across different species.

```
SUN1_HUMAN          VNSALKLYSQDKTGMVDFALESGGGSILSTRCSETYETKTAL-----MSLFGIPLWYFSQ 653
SUN1_MOUSE          VNNALKLYSQDKTGMVDFALESGGGSILSTRCSETYETKTAL-----LSLFGVPLWYFSQ 782
SUN2_HUMAN          VKQALQRYSEDRIGLADYALESGGASVISTRCSETYETKTAL-----LSLFGIPLWYHSQ 586
SUN2_MOUSE          VKQALQRYSEDRIGMVDYALESGGASVISTRCSETYETKTAL-----LSLFGIPLWYHSQ 600
F1R1L9_ZEBRAFISH    VQRALKLYSEDRTGQVDYALESGGGSVLSTRCSETYETKTAL-----MSLFGIPLWYFSQ 855
KOI_DROSOPHILA      VKTVLAIYDADKTGLVDFALESAGGQILSTRCTESYQTKSAQ-----ISVFGIPLWYPTN 832
UNC84_C ELEGANS     IRQMIYEYDTDKTGKVDYALESSGGAVVSTRCSETYKSYTRL-----EKFWDIPIYYFHY 976
SAD1_YEAST          IESTVRKYLTDPVSMPNFALLSTGAEVLPALTSKRYVRRPSAFIPRFTSYFFDSLVVRGH 358


SUN1_HUMAN          SPRVVIQPD---IYPGNCWAFKGSQGYLVVRLSMMIHPAAFTLEHIPKTLSPTGNISSAP 710
SUN1_MOUSE          SPRVVIQPD---IYPGNCWAFKGSQGYLVVRLSMKIYPTTFTMEHIPKTLSPTGNISSAP 839
SUN2_HUMAN          SPRVILQPD---VHPGNCWAFQGPQGFAVVRLSARIRPTAVTLEHVPKALSPNSTISSAP 643
SUN2_MOUSE          SPRVILQPD---VHPGNCWAFQGPQGFAVVRLSARIRPTAVTLEHVPKALSPNSTISSAP 657
F1R1L9_ZEBRAFISH    SPRVVIQPD---MYPGNCWAFKGSQGYLVIRLSLRVIPNGFCLEHIPKSLSPSGNISSAP 912
KOI_DROSOPHILA      TPRVAISPN---VQPGECWAFQGFPGFLVLKLNSLVYVTGFTLEHIPKSLSPTGRIESAP 889
UNC84_C ELEGANS     SPRVVIQRNSKSLFPGECWCFKESRGYIAVELSHFIDVSSISYEHIGSEVAPEGNRSSAP 1036
SAD1_YEAST          EPSIALTPN---NAVAMCWSFQGSEGQLGISLSRPVYVTNVTIEHVQHKIA--HDLSSAP 413


SUN1_HUMAN          KDFAVYGLENEYQ-EEGQLLGQFTYDQDGESLQMFQALKRP--DDTAFQIVELRIFSNWG 767
SUN1_MOUSE          KDFAVYGLETEYQ-EEGQPLGRFTYDQEGDSLQMFHTLERP--D-QAFQIVELRVLSNWG 895
SUN2_HUMAN          KDFAIFGFDEDLQ-QEGTLLGKFTYDQDGEPIQTFHFQAP---TMATYQVVELRILTNWG 699
SUN2_MOUSE          KDFAIFGFDEDLQ-QEGTLLGTFAYDQDGEPIQTFYFQAS---KMATYQVVELRILTNWG 713
F1R1L9_ZEBRAFISH    RRFSVYGLDDEYQ-DEGKLLGDYTYQEDGDSLQNFPVMEEN---DKAFQIIEMRVLSNWG 968
KOI_DROSOPHILA      RNFTVWGLEQEKD-QEPVLFGDYQFEDNGASLQYFAVQNLD--IKRPYEIVELRIETNHG 946
UNC84_C ELEGANS     KGVLVWAYKQIDDLNSRVLIGDYTYDLDGPPLQFFLAKHKP---DFPVKFVELEVTSNYG 1093
SAD1_YEAST          KDFELWVQGMSS--KMFVLLGKARYSLTEDSIQTFSFESSNYIVAEPIQNVILKIKSNWG 471


SUN1_HUMAN          HPEYTCLYRFRVHGEPVK------------------------- 785
SUN1_MOUSE          HPEYTCLYRFRVHGEPIQ------------------------- 913
SUN2_HUMAN          HPEYTCIYRFRVHGEPAH------------------------- 717
SUN2_MOUSE          HPEYTCIYRFRVHGEPAH------------------------- 731
F1R1L9_ZEBRAFISH    HPEYTCLYRFRVHGKPHAQ------------------------ 987
KOI_DROSOPHILA      HPTYTCLYRFRVHGKPPAT------------------------ 965
UNC84_C ELEGANS     AP-FTCLYRLRVHGKVVQV------------------------ 1111
SAD1_YEAST          NPNYTCLYQVRVHGTVPNADEQPIPSLGEKAESTAENTGQDSS 514
```
